# The formation of chromatin domains: a new model

**DOI:** 10.1101/260323

**Authors:** Giorgio Bernardi

**Author notes:** Correspondence and requests for materials should be addressed to G.B. Author Information: The author declares no competing financial interests.

## Abstract

In spite of the recent advances in the field of chromatin architecture^1,2^, the formation mechanism of chromatin domains, TADs, the topologically associating domains, and LADs, the lamina associated domains, is still an open problem. While previous models only dealt with TADs and essentially relied on the architectural proteins CTCF and cohesin, the model presented here concerns both TADs and LADs and is primarily based on the corresponding DNA sequences, the GC-rich and GC-poor isochores, more specifically on their newly discovered 3-D structures. Indeed, the compositionally homogeneous GC-poor isochores were shown to be locally stiff because of the presence of interspersed oligo- Adenines^4,5^, whereas the compositionally heterogeneous GC-rich isochores were found to be peak-shaped and characterized by increasing gradients of GC and of interspersed oligo- Guanines. In LADs, oligo-Adenines induce local nucleosome depletions^4,5^ that are responsible for a wavy structure well adapted for interaction with the lamina. In TADs, the increasing GC levels and increasing oligo-Guanines of the isochore peaks are responsible for a decreasing nucleosome density^5,6^, a decreasing supercoiling^7^ and an increasing accessibility^8^. These factors mould the loops of “primary TADs”, that lack self-interactions since they are CTCF/cohesin-free, yet transcriptionally functional structures^9-11^. This “moulding step” is followed by a second step, in which the cohesin rings bind to the tips of the “primary TADs” and slide down the loops. This process is very likely due to Scc2/Nipbl, an essential factor not only for loading cohesin, but also for stimulating its translocation^12^ and its ATPase activity^13^. This “sliding step” creates self-interactions in the loops and stops at the CTCF binding sites located at the base of the loops that are thus closed and insulated.

---

The human genome (like other mammalian genomes) is compositionally compartmentalized into isochores, large (≥ 0.2 Mb), “fairly homogeneous” DNA sequences that belong to five families, L1, L2, H1, H2, H3, characterized by increasing GC ranges, increasing heterogeneity and increasing gene density^14,15^ (see Supplementary Table S1 and Fig. S1). While the functional importance of isochores was evident for a long time because of their correlations with all the genome properties tested (see Supplementary Table S2 of ref.3) and led to defining them as “a fundamental level of genome organization”^16^, the basic reason for their existence is not yet understood^16^, even if links with chromatin structure and with nucleosome positioning and density were predicted^14,17,18^. Recent investigations^3^ showed, however, that maps of GC-rich and GC-poor isochores of all human and mouse chromosomes match maps of TADs (0.2-2 Mb in size; 1) and maps of LADs (~ 0.5 Mb medium size;2), respectively. Moreover, the average size of human isochores, ~0.9 Mb, is very close to that, ~0.88 Mb, of 91% of the mouse chromatin domains^19^, and both isochores and chromatin domains are evolutionarily conserved in mammals ^17,19^ and correspond to DNA replication units ^20,21^.

The match between isochores and chromatin domains suggested a possible similarity between the 3-D structures of the homogeneous GC-poor and the heterogeneous GC-rich isochores on the one hand and the flat structure of LADs and the loops of TADs, on the other. This working hypothesis was tested by comparing 1) the compositional profile of the DNA sequence of human chromosome 21 (a fair representative of human chromosomes^3,15^) as obtained through non-overlapping 100Kb windows (Fig.1A), and 2) the assembly of the 100Kb sequences into isochores using a sliding-window approach (Fig.1B) with 3) a point-by-point plot of the GC levels of 100Kb DNA segments (Fig.1C).

**Figure 1.**
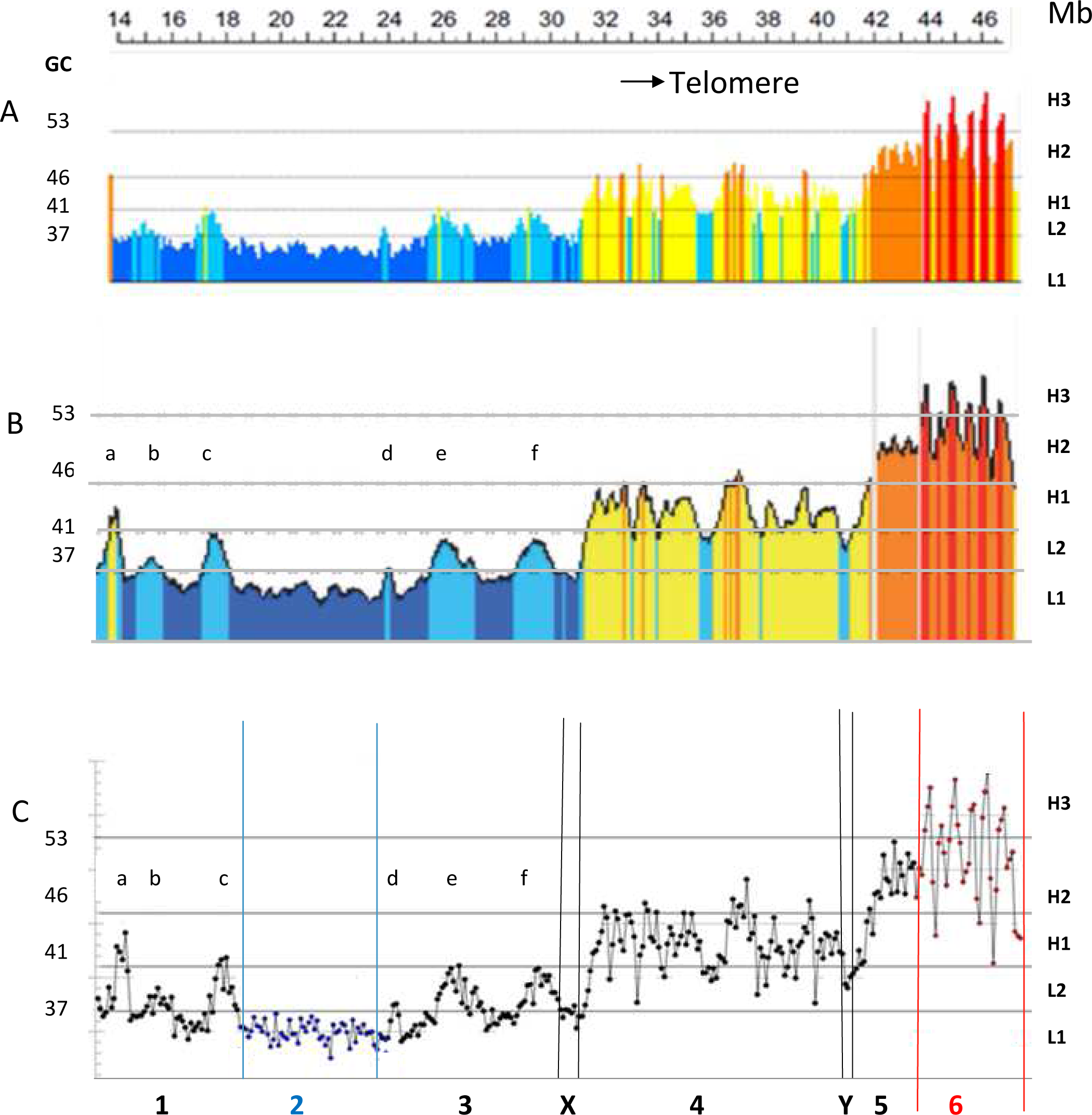
Compositional profile of human chromosome 21. A: Compositional profile through nonover lapping 100Kb windows (from ref.30). DNA sequences from isochore families L1 to H3 are represented in deep blue, light blue, yellow, orange, red, respectively. The left-side ordinate values are the minima GC values between the isochore families (see Supplementary Table S1). **>B**>. **Isochore profile** (from ref. 3) using a sliding window approach. **>C**>. **GC levels of 100Kb segments plotted as points.** Black, blue and red lines, as well as double lines (X,Y), separate regions 1 to 6.

Fig.1C, and its enlarged version Fig.2, showed that isochores can be classified not only compositionally, as done so far, but also topologically, a point missed in Fig.1A for graphical/visual reasons. Indeed, the point-by-point GC profile of chromosome 21 (which was split into several regions) revealed that, while the single peaks (a to f) of regions 1 and 3 and the L1 isochore of region 2 could also be seen in Fig.1 A,B, the isochores from H1 (region 4) and H2 (region 5) families consisted of multiple-peaks emerging from common compositional bases, and the isochores from the H3 family (region 6) were characterized by extremely sharp single peaks ranging from ~45% to 55-60% GC (see Supplementary Table S1). It so happens that chromosome 21 does not comprise homogeneous L2 sequences that are present in other chromosomes (see Supplementary Fig.S2 and ref.3) and that define an L2^−^ subfamily, as distinct from an L2^+^ subfamily formed by the L2 single peaks of regions 1 and 3. Finally, upon close inspection, the peaks of regions 4 to 6 were seen to correspond to the minute peaks of Fig. 1B; this observation allows generalizing the present results to all human chromosomes as investigated by the sliding window approach^3^.

**Figure 2.**
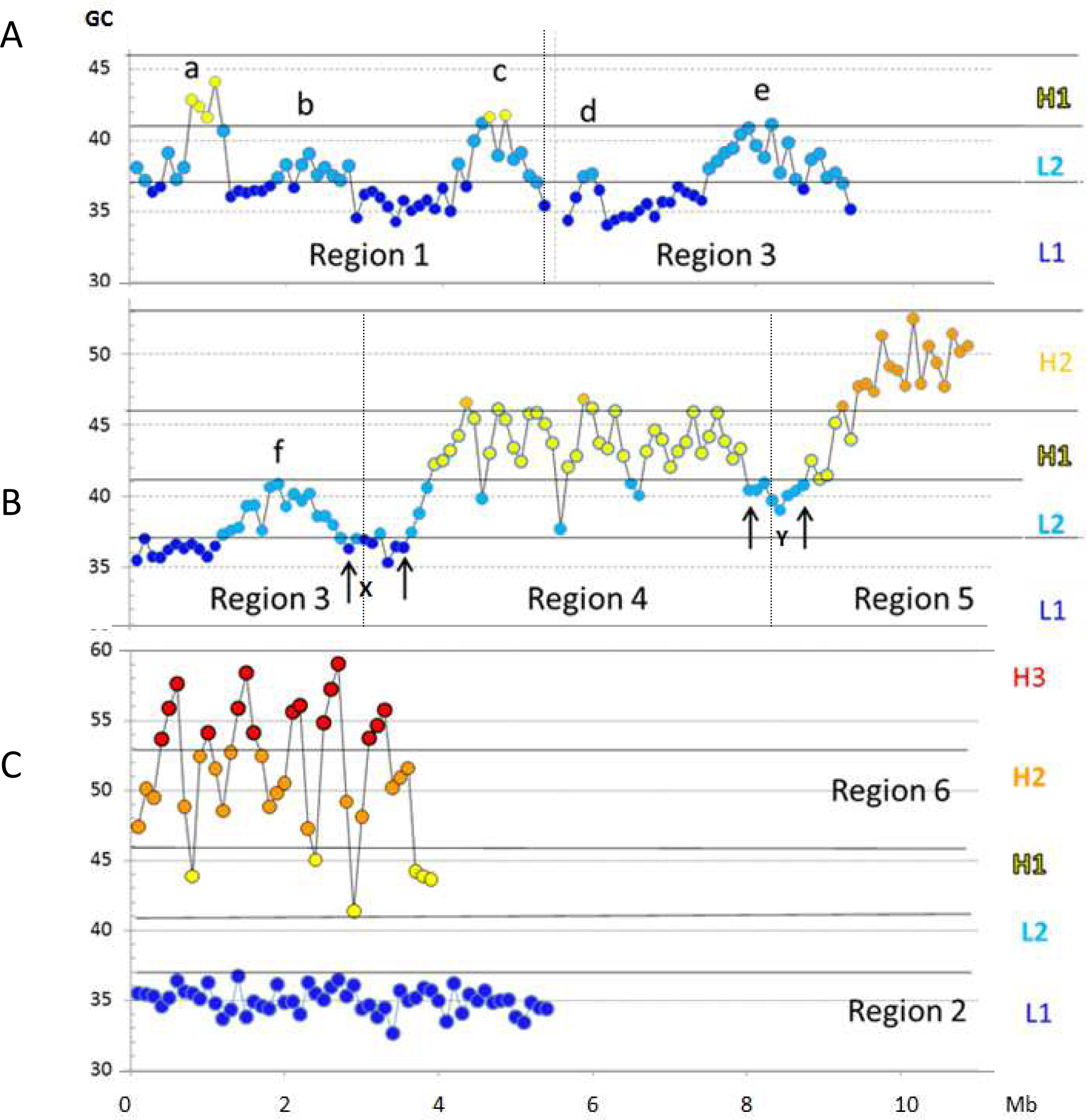
Fig.1C as displayed at a higher magnification. Thick horizontal lines correspond to minima GC values between isochore families (as in Fig.1).

We will now consider in more detail the compositional and topological properties of isochores. In L1 isochores the AT level is ~65% and “A/T-only” trinucleotides represent 25% of the sequences, with AAA/TTT corresponding to 9%^18,22^ and exhibiting a high, fairly constant level which is accompanied by tetra- to octa-A’s (G.Lamolle, V.Sabbia, H.Musto and G.B., paper in preparation). This situation is most likely to be present, even if slightly less pronounced, in L2^−^ isochores since the whole L2 family already comprises 20% of “A/T-only” trinucleotides^18^. In contrast, in L2^+^ to H3 isochores the peaks are characterized by both increasing GC gradients and increasing frequencies of oligo-G’s. In H3 isochores GGG/CCC represent 6% of the sequences^18^, and tetra- to octa-G’s are present in the peaks (while AAA/TTT and tetra- to octa-A’s are present in the troughs between the peaks; Lamolle et al; see above).

Now, one should consider that oligo-A’s and oligo-G’s are intrinsically stiff (for different structural reasons) and strongly inhibitory to nucleosome formation^5^. Two different classes of isochores can, therefore, be distinguished from a topological viewpoint: 1) the L1 (and L2^−^; see also below) isochores that are compositionally homogeneous yet locally stiff and locally nucleosome-depleted^5^; 2) the single-peak L2^+^/H3 isochores and the multi-peak H1/H2 isochores, all these peaks being characterized by increasing densities of GC-rich sequences, interspersed oligo-G’s, CpG’s and CpG islands that disfavor nucleosome formation^5,6^.

Yet another, critical, way to classify isochores is based on their correspondence with chromatin domains. This also leads to visualize the same two groups of isochores. Indeed, the L1 and L2^−^ isochores (the latter including some “valley” isochores such as X and Y of Figs. 1C and 2) correspond to LADs, while the single-peak and the multi-peak isochores of the other families correspond to interLADs (single-loop and multi-loop TADs, respectively; see Supplementary Fig.S2). In the case of L2 isochores, the existence of two sub-families L2^−^ and L2^+^ is confirmed by their correspondence with LADs and interLADs, respectively (see Supplementary Fig.S2).

The results presented so far allow building a model for the formation of LADs and TADs. In the case of LADs (Fig.3A), the presence of interspersed oligo-A’s that causes local DNA stiffness also causes local nucleosome depletions that induce LADs to assume a wavy structure that allows them to adapt and attach to (and even embed in) the lamina^23^.

**Figure 3.**
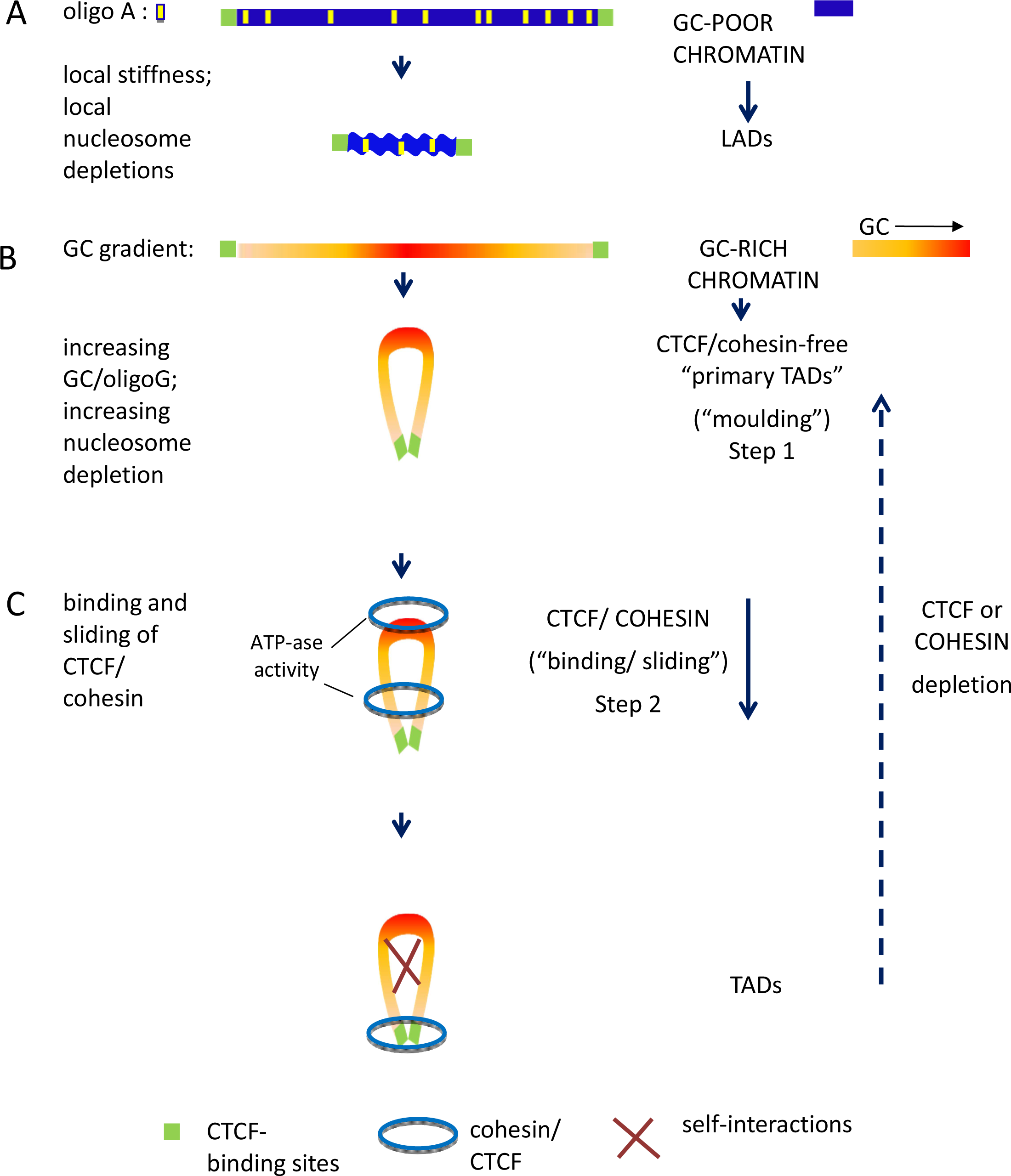
The proposed model for the formation of LADs and TADs. **>A**>. A GC-poor chromatin fibre (blue bar) bounded by CTCF binding sites (green boxes) attaches to the lamina, forming a LAD; the wavy profile indicates the three-dimensional physical adaptation to the lamina by bending and twisting due to interspersed oligo-Adenines (yellow bars). **>B**>. A GC-rich chromatin fibre folds in a “moulding step” to form a single-loop “primary TAD” bounded by CTCF-binding sites (green boxes); the yellow to red color gradient indicates an increasing GC level with increasing frequencies of oligo-Guanines, CpG’s and CpG islands; these features are responsible for the decreasing nucleosome density, decreasing supercoiling and increasing accessibility of the loop; **>C**>. Binding of CTCF and cohesin to the peaks of the loops is followed by a “sliding step” down the loops (helped by the ATP-ase of cohesin sub-units) which ends when the CTCF/cohesin complex reaches the CTCF binding sites at the base of the loops. This process leads to approaching the two branches of the loop, creating self-interactions and closing and insulating the loop. Depletion of CTCF or cohesin^9-11^ reverses TADs into “primary TADs” (broken arrow).

In the case of TADs, their formation appears to take place through a two-step mechanism. In a first step, the increasing GC and increasing oligo-G’s, CpG’s and CpG islands of the isochore peaks are responsible for decreasing nucleosome density^5,6^, decreasing supercoiling^7^ and increasing nuclease accessibility^8^, that constrain chromatin to fold into loops, the tips of the loops corresponding to the highest GC levels. This initial “moulding step”, which involves equal-size chromatin branches (as indicated by the symmetry of GC peaks), leads to CTCF/cohesin-free “primary TADs” (Fig.3B), that are already functional since CTCF or cohesin depletions are not followed by any remarkable alteration of transcription as recently shown^9-11^. Interestingly, a wider nucleosome spacing and higher flexibility of the loops compared to boundary regions was also proposed by the “insulation/attraction model”^1^ (see Supplementary Fig.S3).

Two points should now be raised about the “moulding step”. The first one concerns which one of the two factors, the 3-D structure of isochores or the nucleosome density, is primarily responsible for moulding; the demonstration that TADs are not very different in fibroblasts and in spermatozoa^24^ (in which case nucleosomes are replaced by protamines) provides an answer in favor of the first possibility. The second point is that the existence of “primary TADs” solves the notorious problem of the energy source required by the active extrusion model^25,26^, which involves pulling megabase-size chromatin fibres (with branches of different lenghts because of the random location proposed for the extrusion starting points) through the CTCF/cohesin ring. Indeed, while in the case of condensin an ATP hydrolysis dependent molecular motor was recently identified, this was not the case so far for cohesin^27^ (see, however, below).

In a second step (Fig. 3C), one may consider that the CTCF/cohesin ring complex binds to the peaks of the loops and starts a “sliding step” down the “primary TADs”. In this connection, it is of great interest to consider the recent proposal^12^ that Scc2/Nipbl, an essential factor for loading cohesin on the loops, may also stimulate cohesin’s translocation along the loops. The ATPase activity associated with cohesin sub-units Smc1 and Smc3 ^13^, also stimulated by Scc2/Nipbl would provide the energy necessary for sliding, an energy obviously far below that required for the active extrusion (see above). This sliding step leads to closer contacts between the two branches of the loops and to self-interactions and stops at the CTCF binding sites, that are located at the base of the loops, thus closing and insulating them. Incidentally, the possibility that extrusion could also be mediated by a single cohesin ring sliding over the top of a preformed chromatin loop was already taken into consideration previously^28^.

In conclusion, the two-step “moulding/sliding” model of chromatin domains of Fig.3 represents a paradigm shift compared to previous models (presented in Supplementary Fig. S3) that essentially rested on architectural proteins and neglected, as also remarked elsewhere^29^, the role played by DNA. Indeed, the new model, which provides the same basic explanation for both TADs and LADs, essentially relies on well-established physico-chemical properties of DNA sequences, on the newly discovered 3-D topology of isochores and on its connection with chromatin domain architecture.

This investigation, which only dealt with the evolutionarily conserved features of chromatin architecture, also leads to three general conclusions: 1) isochores are not based anymore only on their compositional properties and on their correlations with all genome properties tested^3^, but also on their essential function of encoding their own 3-D topology, the primary factor in moulding TADs and LADs; 2) the definition of “genomic code”, originally proposed^14^ for the set of general rules concerning isochores and for properties associated with isochores, should now indicate the basic phenomenon, namely the encoding of the 3-D isochore structures and, as a consequence, of chromatin domains; 3) non-coding sequences, that represent ~98% of the human genome, play a major role in defining chromatin architecture; this provides yet another argument against the decades-old “junk DNA” hypothesis. Finally, this investigation raises the question of the possible functional differences linked to the different isochore structures of LADs (L1 vs L2^−^) and TADs.

## Acknowledgements

The author thanks Paolo Ascenzi for hospitality, Giacomo Bernardi (Santa Cruz, California, USA), Oliver Clay (Medellin, Colombia), Paolo Cozzi (Milan, Italy) and, especially, Kamel Jabbari (Cologne, Germany) for critical reading, comments and discussions. Caterina Nuvoli and Marta Ritucci provided excellent technical help. This research was supported by the Kimura Prize for Molecular Evolution and Evolutionary Genomics conferred to the author (Tokyo, June 2016).

